# Proprioception impacts body perception in healthy aging – insights from a Psychophysical and Computational Approach

**DOI:** 10.1101/2024.07.23.604821

**Authors:** G. Risso, M. Bieri, T. Bertoni, G. Mastria, L. Allet, A. Serino, M. Bassolino

## Abstract

The experience of owning a body (body ownership, BO) and the perception of our body dimensions (metric body representation, mBR) depend on the integration of multisensory cues. As the human sensory system is subjected to a decline along the lifespan, encompassing all sensory modalities, we hypothesize that body perception may be different in older, as compared to young adults. Here, we investigate this hypothesis by comparing the multisensory processing underlying BO and mBR in healthy older (> 65 years) and young individuals. First, we applied rigorous computational and psychophysical methods to assess alterations in mBR and BO quantitatively. We then modeled the manifold relationship between the observed body misperceptions and the potential underlying sensory, motor, and cognitive factors. The results highlight significant differences between the two groups, with higher distortions in perceived arm dimensions and an increased tendency to experience BO towards a virtual hand in the aged group. These differences in both mBR and BO are explained by the reduced proprioceptive abilities of older adults, suggesting a crucial role of proprioception in driving age-dependent plasticity in body representations. Overall, our modeling and experimental approach provide new perspectives on altered body perception during aging, suggesting that they stem from the physiological proprioceptive decline occurring in older adults, and laying the groundwork to generate prevention and stimulation strategies to restore accurate body perception in aging.

## Introduction

The human sensory system is subjected to a decline along the lifespan^1^, encompassing all sensory modalities^1^, i.e. vision^2^, hearing^3^, taste, smell^4^, tactile and proprioceptive sensitivity^5, 6^. Moreover, older adults suffer from reduced functionality of the motor system, with decreased stability^7, 8^ and balance control^9^, increased risk of falls^10, 11^, and motor execution time^12, 13^. Changes during aging concern also how these different modalities are integrated, (see^14, 15^ for a review). Coherent multisensory integration of bodily cues during daily life interactions is necessary to build different brain representations of the body, underlying perception, action, and self-consciousness^16–18^. Among those, a metric body representation (mBR^16, 19^) is built and continuously updated via the online integration of visual, proprioceptive, and tactile cues to implicitly code the size and shape of different body parts, which is crucial for correct motor control^20, 21^. Furthermore, the natural experience of the body as our own, i.e. body ownership (BO), also depends on the coherent integration of bodily cues in space and time. This has been demonstrated by alteration in BO induced by the so-called multisensory body illusions (e.g., the rubber hand illusion, RHI^19, 22^) or by alteration in the processing of somatosensory and in particular proprioceptive information as in amputees^23–26^ or stroke patients^27–31^ reporting altered body experience e.g. phantom or altered perceived dimension of the affected limb, mBR.

Therefore, and considering the well-documented deterioration of sensorimotor processes, including proprioceptive ones^5, 6^, during physiological aging^14, 15^, an emerging view suggests that possible changes in BO and mBR can occur also in healthy older adults^14, 32–34^. However, evidence in this direction is scarce.

Previous studies investigating age-related changes in BO applied the RHI^35^. In this paradigm, the illusion that a rubber hand belongs to the participants’ body is induced by manipulating the congruency between visuo-tactile stimulation of the participants’ real limb (hidden from sight) and a rubber hand^22^. Explicit ownership ratings towards the rubber hand and its perceived position are reported to be not different in seniors rather than young adults in some studies^36, 37^, significantly decreased (i.e. lower illusion)^32, 33^ or significantly increased (i.e. higher illusion) during aging in others^32, 33^. Thus, whether and how BO is affected during aging is still a matter of debate. In addition, the origin of these age-related changes, if any, has not been systematically investigated.

Related to the mBR, only two seminal studies are available, both suggesting changes during aging with older adults perceiving their arm as shorter than younger controls^34, 38^. While these results are consistent, the sensorimotor or cognitive factors driving such distortions remain unknown.

Thus, there is limited, and not conclusive evidence related to the alteration of body representation, and the potential associated factors, in healthy older adults. Bodily awareness is a central component of human sensation, cognition, and action. Without accurate knowledge about body perception during aging, we also lack potential strategies to counteract alterations in this function and its related factors, with an impact on quality of life and autonomy^14^. Accordingly, this work aims to (i) quantify age-related alterations in mBR and BO, (ii) as well as the sensorimotor and cognitive factors underlying those distortions.

To these aims, we applied the same task previously used ^34^ to assess mBR in older adults, but with the addition of concomitant assessments of sensorimotor and cognitive functions putatively declining with age. Furthermore, to overcome the controversial results reported by previous RHI-based studies on BO in aging^32, 33, 36, 37^ we adopted an alternative recently proposed approach allowing to quantify BO with unidirectional predictions and to model the contribution of uni- and multi-sensory processing to ownership^39–42^. In this paradigm, participants perform a virtual reality-based reaching task towards visual targets, while observing a virtual hand displaced with a variable visual angle on a trial-by-trial basis with respect to the actual position of their real hidden hand^43^. The reaching errors induced by the incongruent virtual hand and the subjective ratings of ownership for the virtual hand are respectively considered implicit and explicit indexes of BO^39^, allowing to overcome the limitations of task-based purely on explicit questionnaires. These behavioral indexes are analyzed in a Bayesian causal inference framework, in which the likelihood of a limb belonging to one’s body is estimated based on the integration of multisensory body information, depending on the precision of visual and proprioceptive signals. Thus, this approach allows to quantify age-related alterations in BO and to model the underlying uni- /multi-sensory and cognitive components.

In line with previous studies documenting alterations in body representations in individuals with proprioceptive impairments^23, 26, 31^, and considering the physiological decline of proprioception during aging ^5, 6^, we expect i) to identify higher mBR distortions and altered BO in older adults and ii) to observe a predominant role of proprioception in explaining these alterations.

## Results

### Older adults show a higher tendency to experience ownership towards a virtual hand

To measure age-related alterations in BO quantitatively, we applied the visuo-proprioceptive disparity (VPD) task (see Figure 1A). 23 healthy older individuals (HO; 8 males; mean age: 73.82 ± 6.36) and 23 healthy young adults (HY; 12 males; mean age: 27.17 ± 4.27) participated to the experiment.

**Figure 1.**
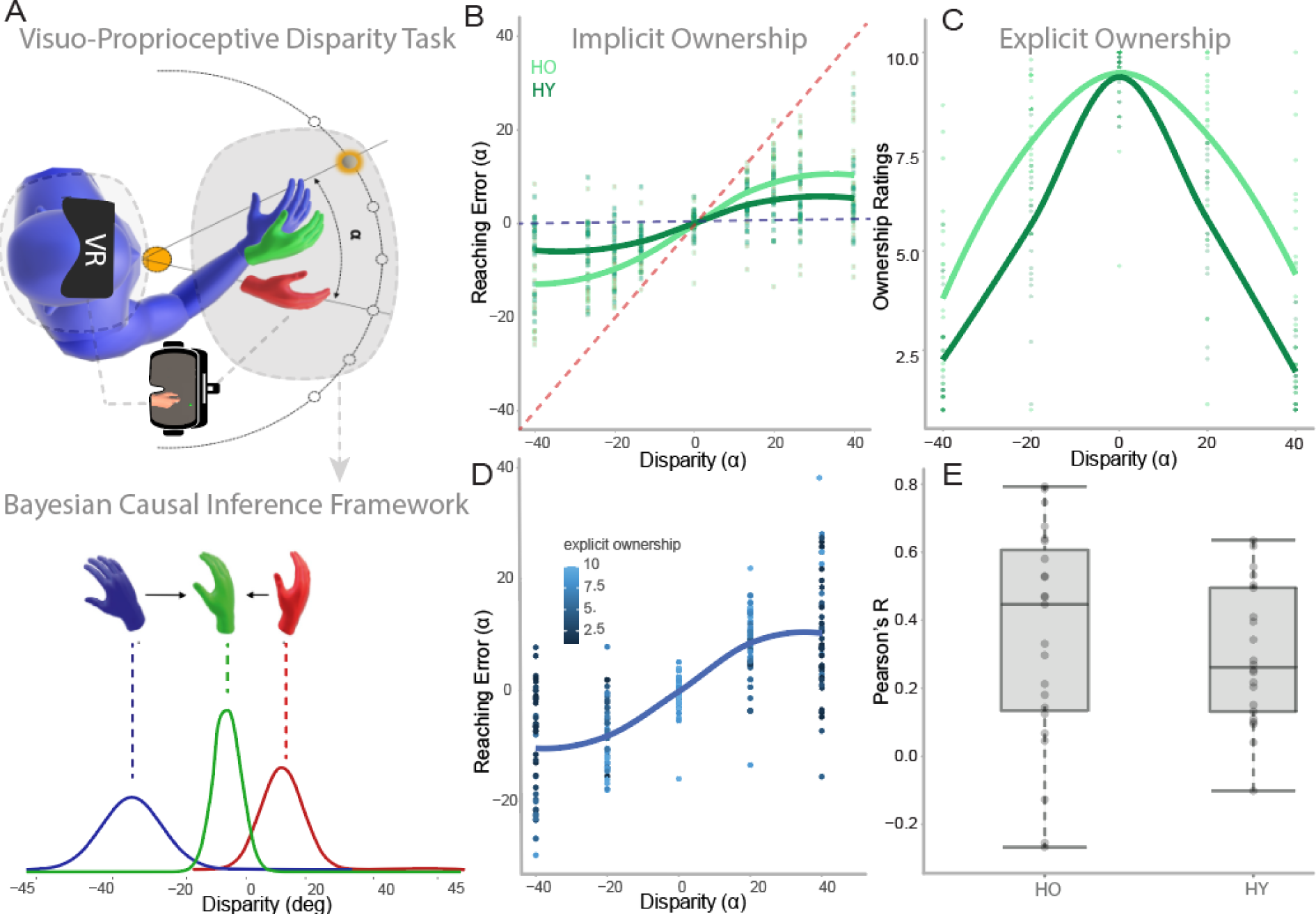
**A) Experimental setup**: The Visual Proprioceptive Disparity task (VPD) consists of a reaching task in which at each trial participants are required to perform a reaching movement from the starting position to one of the seven possible target positions. The participants are immersed in a virtual reality environment where they observe a virtual hand (in red) which in different trials, could either be in a position congruent (0 disparity) with the real hand (in blue and hidden from the view) or in an incongruent position at a certain angle α of disparity (7 possible angles from α = - 40° to α = ± 40°; where positive disparities correspond to a clockwise rotation of the virtual hand with respect to the real one, while counterclockwise rotation is provided with negative disparities). The participants’ task is to reach the target by following their real hand as closely as possible. On each trial, the difference between the real hand (in blue) and the movement made by the participant (in green) is calculated (reaching error). No feedback about the errors is provided to avoid corrections. VPD data was fitted with the Bayesian causal inference model (Bayesian CI). Panel A bottom row shows the hand position estimates according to the Bayesian CI model as a function of spatial disparities between the visual-virtual (red) and proprioceptive-real (blue) hands. According to the Bayesian CI framework, the final reached position (in green) will be a weighted average between pure visual and pure proprioceptive guidance. The relative weight of vision and proprioception will depend on their relative precision and on the a priori probability computed by the brain that the virtual hand is one’s own. **B) Older adults (HO) show higher implicit ownership towards the virtual hand than younger adults (HY).** The plot shows the mean values of the Reaching error of each participant for each level of disparity between the real and virtual hand. The horizontal blue dotted line at zero error represents the participants’ performance if they consider only the proprioceptive signal (i.e. if they reach the target following their true hand in blue). The red diagonal dotted line represents the participants’ performance if they follow only the visual signal (i.e. if they reach the target following the virtual hand in red). Older adults’ (in dark green) performance is closer to the red diagonal line thus suggesting that they weigh the visual signal more and tend to follow the virtual hand more than young adults. **C) Older adults show higher explicit ownership towards the virtual hand than younger adults.** The plot shows the mean ratings reported by each participant for each level of disparity between the real and virtual hand. In both groups, as expected, higher levels of ownership towards the virtual hand are reported for lower visuo-proprioceptive disparities. However, older adults (dark green) reported a higher level of ownership toward the virtual hand for each level of disparity. **D**) **Relationship between explicit judgments and implicit behavior.** The plot shows the mean values of Reaching Error of each participant (HO and HY) in the third and final block, during which participants also reported ownership ratings for each trial. As expected, at lower disparities, higher ownership (represented by light blue dots towards 10, in the legend) was reported. Importantly, for higher disparities, participants showing higher reaching errors (i.e. following more the virtual hand) reported also higher ownership ratings suggesting a relationship between the implicit and explicit ownership experience. **E) Explicit and implicit ownership correlations in young and older adults.** The plot shows individual Pearson’s correlation coefficients between residual reaching errors and ownership ratings calculated trial-by-trial in the two groups.

In the VPD task, participants were asked to reach a visual target while seeing a virtual hand displaced with respect to the actual position of their real, and not visible hand, with 7 possible displacement angles, varying on each trial. Participants were asked to reach following their real hand (proprioceptive cue). In the last of the three blocks (see methods), they were also asked to report their agreement with the statement “I felt as if the virtual hand was my hand” on a Likert scale from 1 to 10 (maximum agreement) on each trial, for the different disparity levels (see Methods).

As visible from Figure 1C, Healthy Older adults (HO) reported significantly higher explicit ownership ratings than the younger ones (HY; Linear Mixed Model (LMM): β = -1.59 | 95% CI [-2.35, -0.82], group: t(1603) = -4.08, p < .001; Std. beta = -0.46, 95% CI [-0.67, -0.24]. Also, the reaching error (Figure 1B; defined as the distance between the target position and the real hand position at the end of the reaching, see Methods for the details) was higher (i.e. towards the virtual hand) in older adults than younger adults, and the difference was particularly significant for negative disparities (i.e. when the virtual hand was displaced with clockwise rotation with respect to the real one, LMM: beta = -0.16, 95% CI [-0.18, -0.15], t(6034) = -19.70, p < .001; Std. beta = -0.36, 95% CI [-0.40, -0.33]). As evident in Figure 1B the reaching movements of older adults are more shifted towards the virtual hand (i.e. towards the red dotted diagonal line) than the movement performed by the younger ones. Interestingly, the reaching error covaried with the explicit ownership ratings at a trial-by-trial level (see Figure 1E), suggesting that the reaching error observed in the behavioural task is effective in capturing implicitly the experience of ownership perceived by the participants (see Figure 1D and E).

### Older adults show more bias in metric body representation

To investigate alterations in mBR concerning the upper limb, we applied the Body Landmarks Localization task (BL; see Figure 2A), previously used in several studies to investigate the perceived dimensions of participants’ body parts^21, 34, 44^. The same 23 healthy older individuals involved in the VPD experiment and other 23 healthy young adults (HY; 8 male, mean age: 24.82 ± 4.09) participated in the experiment. Participants were asked to localize specific landmarks on the upper limb while their arm was hidden from their view. The perceived lengths of the different body parts were indirectly computed as the distance between the different landmarks, and the main output for the analyses consists of an index of estimated distance ^21^: 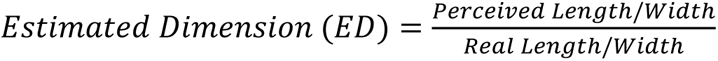 Older adults significantly underestimated their arm length as compared to younger adults (Wilcoxon signed-rank: z= -2.24, p=0.025; see Figure 2B left panel). No other significant comparisons emerged see Figure 2C and B right panel).

**Figure 2.**
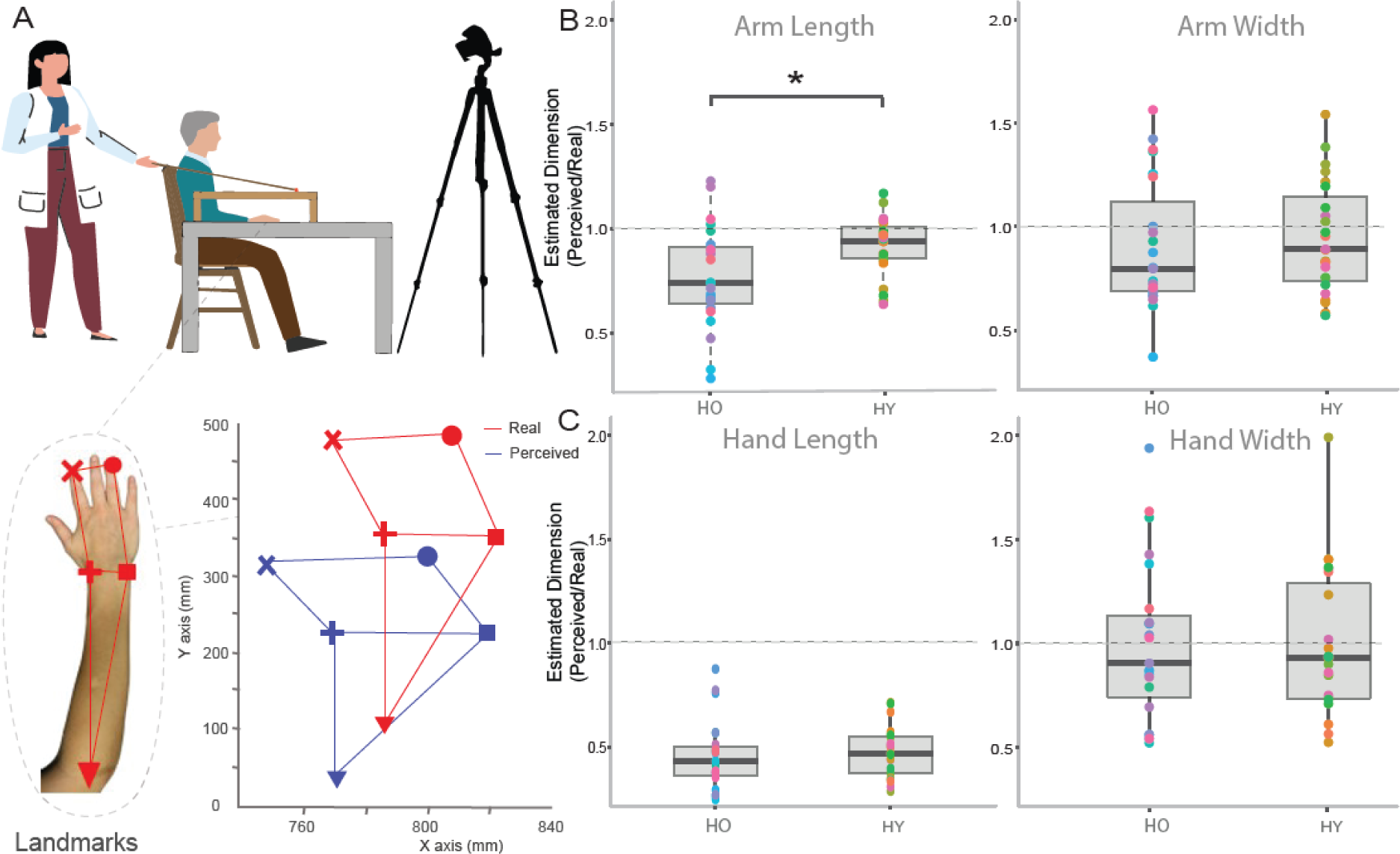
**A) Body Landmarks Localization Task.** The top panel shows the experimental environment in which the participant is seated with the arm resting in a fixed position hidden from the view by a table. At each trial, the participant guides the experimenter holding a stick with a marker on the perceived location. The localization is recorded thanks to a tracking system. The five landmarks that the participants were required to localize (tip of the index, tip of the ring, internal and external wrist, and elbow) are shown in the bottom row, together with a possible example of the offline reconstruction of the tracked localization of the real landmarks (in red) and the perceived landmarks (in blue). **B) Older adults underestimate arm’s length with respect to younger adults.** The panel represents the estimated width and length (i.e., the ratio between the perceived and the real size) of the hand. Values below 1 (dashed line) indicate an underestimation, while values above 1 indicate an overestimation. While no differences can be observed for the arm width perception, older adults significantly underestimate the arm length with respect to younger ones(*p<.05); **C) Older and younger adults show no differences in hand estimated dimensions**. No differences in the hand estimated perception (neither for the length nor width) can be observed between the two groups.

### Factors underlying the age-related distortions in body perception: a fundamental role of proprioception

To better understand the sensorimotor and cognitive components at the origin of the BO alterations in older adults assessed with the VPD task, we took advantage of the Bayesian causal inference framework. Accordingly, the best way to interpret a stimulus is by performing a weighted estimation between the sensory cues, where the weights should be proportional to the reliability of the sensory cues ^39, 45^, so that more reliable sensory modalities contribute more to the final estimate. This framework can be readily transposed to the VPD task. Indeed, to reach the target with their real hand, the participants combine spatial information from proprioception about the real hand position with (incongruent) visual information about the virtual hand position, and the final reached position is a weighted average between proprioceptive-based and visual-based guidance. The key assumption of our model is that the more ownership is attributed to the virtual hand, the more weight will be attributed to visual cues in performing the reaching movement ^39, 40^. The processes to integrate the sensory information, i.e. the relative weight attributed by the participants to vision (virtual hand) and proprioception (real hand), will depend i) on one side on “bottom-up” *sensory* components, depending on their relative precision. If proprioception is less reliable than vision, participants will rely more on visual information about the virtual hand during reaching (e.g. if proprioception is more reliable than vision, the higher weight will be given to the proprioceptive signals); ii) on the other side, it will depend on a top-down *cognitive* component consisting in the a priori probability that the visual information from the virtual hand and the somatosensory information from the real hand originate from a unique object (that is, they have a common cause), i.e. the a priori probability that the virtual hand is one’s own. A higher prior will lead to higher ownership values, and thus to a higher weight of visual cues in guiding reaching movements (see Figure 1A).

In this view, we modeled our task as the result of a Bayesian causal inference process and the sensory and the cognitive components are extracted from the Bayesian Causal Inference model and fitted from the multisensory task^40, 43^, as follows: i) the sensory components computed as the weight given to each sensory cues, i.e. the sensory inputs precision (respectively, visual: σ_v_ and proprioceptive: σ_p_); ii) the cognitive component computed as the prior of a common cause P_comPrior_ among the visual and sensory signals.

Related to the sensory component, and as visible from Figure 3A, the proprioceptive (σ_p_) [Wilcoxon Signed-Ranks Test: z = -3.797, p<.001], and visual (σ_v_) [z = -2.257, p<.05] variability of the older adults at the VPD is significantly higher than that of the younger adults, indicating that older adults have significantly higher sensory variability in their real hand localization. However, in older adults σ_p_ is significantly higher than σ_v_, meaning that for older adults’ vision is the most reliable sensory cue during the VPD task [z = -1.973, p<.05]. Conversely, in younger participants, the difference between σ_p_ and σ_v_ shows the opposite direction with proprioception being the most reliable sensory cue [z = -3.221, p=.001], following the experiment instruction to follow the real hand. To confirm that the older adults’ results related to proprioceptive precision extracted from the model truly reflect their proprioceptive accuracy, we used an independent, purely unisensory, proprioceptive task^39, 42^ i.e. the static proprioceptive judgment task (PJ; see also Table 2 in the Methods). Our results confirm the prediction of the model: the proprioceptive precision observed at the PJ (σ_PJ_) and the one extracted from the model (σ_p_) correlate [r(44) = 0.554, p < .001; see Figure 3C]. This result shows that the multisensory behaviour (from which σ_p_ was extracted) truly reflects the sensory variability observed during a unisensory proprioceptive task (σ_PJ_). Furthermore, in line with the results in proprioceptive precision (σ_p_) extracted from the VPD (figure 3A), our results show that also at the unisensory PJ task, older adults show significantly lower proprioceptive precision with respect to the younger participants [z = - 5.416, p<.001; see Figure 3B]. Finally, to verify whether a decreased proprioceptive precision is also linked with the explicit experience of ownership towards the virtual hand, we correlated the mean explicit ownership ratings with the PJ proprioceptive variability. There was a significant positive correlation [r(44) = 0.477, p < .001; see figure 3D], confirming that participants with a lower proprioception precision, also report a higher level of explicit ownership towards the virtual hand, thus further confirming the relation between a lower proprioceptive precision and the tendency to embody an external object as part of their own body.

**Figure 3.**
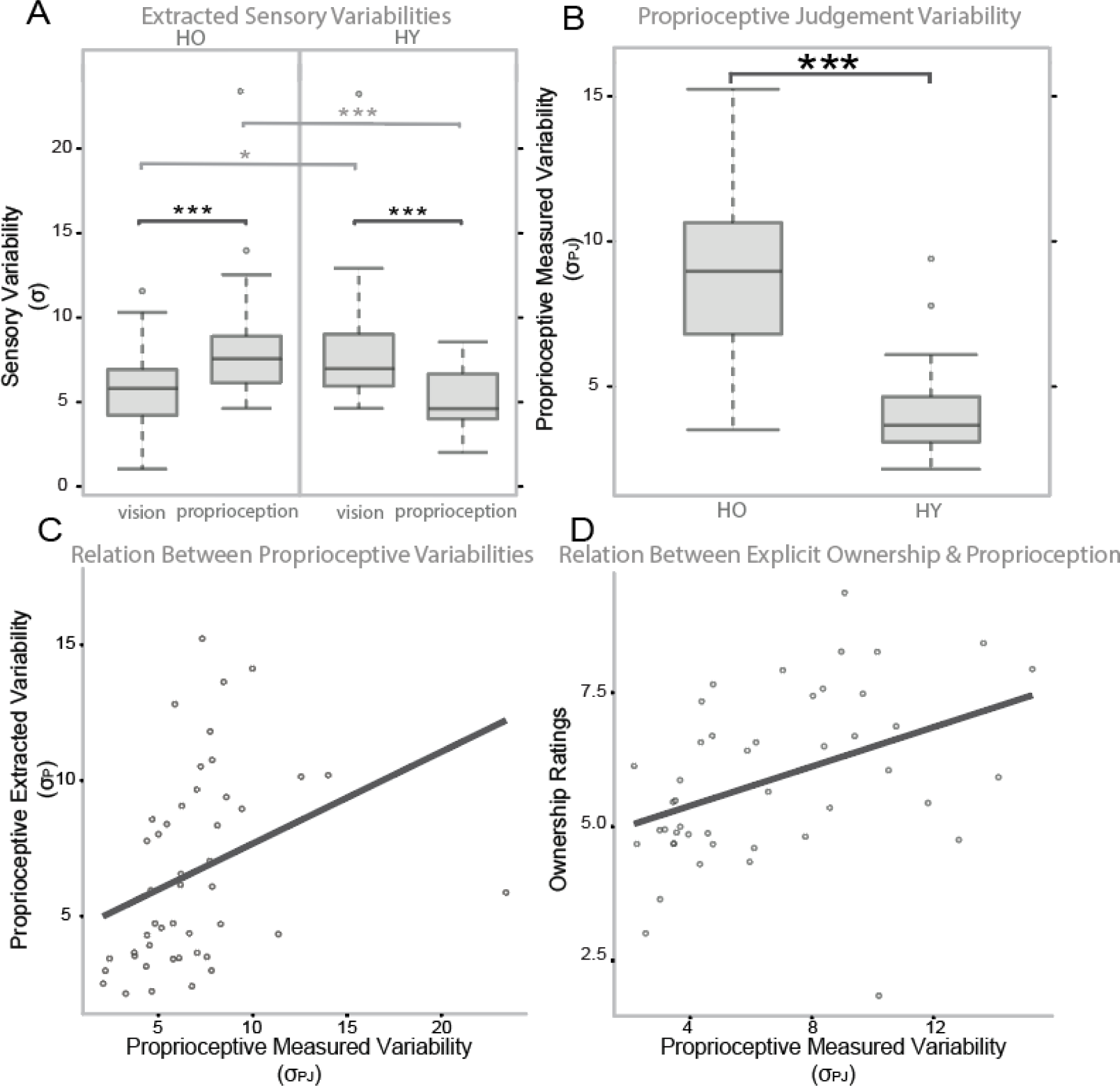
**A) Visual dominance at reaching VPD task for the older adults but not for the younger ones:** The visual (σ_V_) and proprioceptive (σ_P_) variabilities of the participants extracted by the model from the multisensory reaching performance to the VPD task are shown for older adults (HO) and younger ones (HY). Higher σ values represent higher variabilities. The asterisks indicate significant differences between groups (in light grey) and within group (in black): ***p<.001; **p<.01; *p<.05. **B) Younger adults show higher proprioceptive accuracy than older adults:** The figure shows the comparison between the proprioceptive variability σ_P_ of the two groups directly measured through a purposely designed task (PJ). **C) Linear relationship between the proprioceptive performance during the multisensory reaching task (**σ_P_ extracted from the VPD task) and the variability directly assessed with the **unisensory proprioceptive task** (σ_PJ_ assessed with the PJ task). **D) Linear relationship between the explicit judgment of ownership** (trial-by-trial mean ownership ratings) **and the proprioceptive performance** (σ_PJ_, variability at the PJ task).

Related to the cognitive component, we found that the P_comPrior_ is not significantly different between the two groups [z = -0.847, p=.397]. This result suggests that the observed effect of aging in the way visuo-proprioceptive bodily signals are integrated into a unique object of perception can be attributed to a different precision of unisensory processing and not multisensory integration priors.

Taken together, these findings related to the ownership experience assessed through the VPD task suggest that increased proprioceptive variability, and not changes in cognitive priors, drive the increased tendency to embody a virtual hand in older adults compared to younger ones.

There is no validated computational model to interpret the results of the mBR task. Therefore, to investigate the factors affecting the dimensional perceptual bias observed in older adults, we administered to the older participants an independent battery of tasks assessing functions that might have a role in mBR. mBR is built up and updated through the continuous flow of sensory information about our own body from various sensory ^44, 46^ and motor ^20, 21^ sources, as well as it is influenced by higher-level cognitive factors ^16^. To assess the different components playing a relevant role in affecting mBR during aging, we selected tasks tapping into these factors, with a preference for those already applied in aging ^47–50^. For the sensory components, we evaluated tactile, visual, and proprioceptive cues (assessed respectively through two-point discrimination, proprioceptive judgment, and open loop, see Methods, Supplementary Materials, and Table 2 for more details). For the motor component, we included subjective and self-reported assessments of participants’ daily lives and direct measurements of movement and strength (physical activity ratings, speed of walk, max arm grip strength evaluation with Jamar). Moreover given the impact of cognitive flexibility^51^, short and working memory^49^ decline in aging, we added tasks assessing these executive functions (the digit span forward and backward, trial making test; see Table 2 and Methods for further details).^48^ As a unidirectional measure of distortion in mBR, we calculated an index of bias by computing the Estimated Error Index defined as follows: *Estimated Error* (*EE*) = |*ED* - 1|. The lowest possible value is 0, i.e., no bias in the arm length perceived dimension, while higher EE values represent higher errors. Then, to identify the set of factors explaining the observed distortions, we applied a stepwise linear regression on EE values by using the outputs of the tasks assessing sensory, motor, and cognitive functions as predictors.

The best model predicting the arm length Estimated Error (EE) is reported in Table 1. It significantly fitted the data explaining the 56% of the variability of the model (43% R-squared adjusted; F(5,17)=4.277, p=0.011, see methods for more details on the variable selection) and included: i) the level of physical activity, ii) the isometric arm grip strength iii) the forward and iv) backward memory span, and v) the response variability of the OL task assessing the proprioceptive ability in dynamic condition. Among those, the parameters significantly predicting the arm length bias EE in older participants were the isometric handgrip strength (β = -0.014, p = .026), the backward memory span (β = -0.182, p = .002), and the response variability of the proprioceptive dynamic task (β = 0.020, p = .030). Taken together these results suggest that specific proprioceptive, motor, and cognitive factors underlie the metric bias on the arm observed in the older adults.

**Table 1:** Stepwise Linear Regression Output. The table represents the output of the stepwise linear regression having the Estimated Error on the arm length as the dependent variable. The first column indicates the sensory, motor, and cognitive potential predictors of bias in mBR and considered in the analyses. The second column summarizes the task used to evaluate each factor. The third column indicates whether the variables resulted as explanatory variables according to the Stepwise regression method to predict the Estimated Error (EE) on the arm. The fourth and fifth columns report, respectively, the coefficients and the associated p-value of the resulting linear regression model (significant predictors, p<.05, are highlighted in bold).

| Variable | Task | Stepwise Included | Regression Coefficient | Regression p-value |
| --- | --- | --- | --- | --- |
| Arm Tactile Acuity | Two Points Discrimination task | No | / | / |
| Proprioceptive variability in static condition | Proprioceptive Judgment task | No | / | / |
| Proprioceptive variability dynamic condition | Open Loop task | Yes | 0.02 | **0.029** |
| Level of physical activity | Questionnaire: ordinal scale with 6 levels | Yes | 0.07 | 0.089 |
| Speed of walk | 4-meter walk | No | / | / |
| Isometric arm grip strength | JAMAR hydraulic dynamometer | Yes | -0.014 | **0.026** |
| Working memory | Digit Span Backward | Yes | -0.18 | **0.002** |
| Short-term memory | Digit Span Forward | Yes | 0.05 | 0.072 |
| Cognitive Flexibility | Trail Making Test B (time of execution) | No | / | / |

## Discussion

The present results highlight significant differences between young and older adults in body perception, including the two studied components, namely Body Ownership and metric Body Representation.

First, differently to some previous studies exploiting the RHI^32, 33^, our findings reveal a similar higher explicit (i.e. when participants were directly asked to make a judgment) and implicit (i.e. the error observed in the behavioral reaching task) ownership towards a virtual hand in older as compared to young adults. With respect to previous works based on RHI, the current approach has not only the advantage of providing a great amount of quantitative data (reaching error computed on 144 trials and ownership judgment collected on 35 trails versus typical few proprioceptive judgments and twice a few-item questionnaire collected after sessions of synchronous and asynchronous stimulation) but also to parallel testing and modelling the contribution of the different sensory and cognitive components to BO. Our results shed light on an aspect that has long been the subject of debate^14^, i.e. whether any bias in the attribution of ownership to a fake hand in aging is due to (i) higher-level cognitive components (i.e. priors, e.g. due to learning, experience, suggestion ability or demand characteristics ^14, 33, 37, 52, 53^), or (ii) changes in sensory abilities that may occur with age^14, 15, 36, 54, 55^. Our results show a significantly diminished ability to rely on proprioceptive cues during reaching in the elderly and no significant difference in their priors clearly supporting the second hypothesis. Thus, a higher tendency to attribute ownership to a virtual hand, even when visually displaced from their own hand, shown by the older adults can be explained by their lower proprioceptive sensitivity with respect to younger controls.

This conclusion is confirmed also by the explicit ownership ratings, whereby older adults demonstrate a higher rating of embodiment towards the virtual hand at all the disparities, and this effect correlated with reduced proprioceptive precision. To summarize, older adults show a wider spatial window in integrating visual information about the virtual hand with proprioceptive information about their real hand at higher disparities.

Interestingly, altogether these findings are in line with results observed in young participants and previous studies exploiting this same paradigm^39^ where normal fluctuations towards poorer precision in proprioception indicate a higher tendency to attribute the virtual hand as they own ^39^. Importantly, the present results in older adults are in line with those recently obtained in post-stroke patients with proprioceptive impairments. By applying the same experimental and computational approach, results demonstrated that patients showed a higher attribution of a virtual hand to their body both at implicit (reaching error) and explicit (ownership ratings) levels, specifically for the limb contralateral to the affected hemisphere. Importantly, the alterations reported in stroke patients on their affected side were more evident than those found here in healthy aging, as evident from the comparisons with an age-matched control group. Interestingly, the worst results in stroke patients were not explained solely by proprioceptive deficits, but rather by patients’ lesions of the frontoparietal network, suggesting that, in brain-damaged patients, BO alteration is caused by two parallel mechanisms, i.e., a reduction of proprioceptive precision and an impairment of multisensory integration functions due to damage to the frontoparietal network. In this previous work on stroke patients, we also found that the increased tendency to embody an incongruent virtual hand of the patients at the VPD task was related to the disembodiment of patients’ real hand ^42^. In other words, patients not only reported higher implicit and explicit ownership towards the virtual hand when there was a disparity with their real hand, but also, they reported less ownership towards the virtual hand when it was congruent with the real hand (i.e. at zero disparity). In the current study, we found no evidence of disownership for the own body in the older adults (see also Supplementary Materials). Together, the present findings from healthy older adults and the previous findings from stroke patients agree in pointing to a fundamental role of proprioception in the modulation of BO in healthy and pathological conditions, with additional intervening factors in patients concerning damage to frontoparietal network and disownership of one’s own limb.

Furthermore, we found an altered mBR in older adults who perceived their arms as significantly shorter than younger controls. These results replicate previous findings obtained in another sample of aged participants collected with the same task by our group ^34^ and with another approach (i.e., the arm bisection task) by other authors ^38^. However, no previous work investigated any possible factors underlying these modifications in mBR. Distorted characteristics in the perceived size of body parts are common also in healthy young populations (in particular related to hand size) and it has been replicated under different tasks and conditions ^21, 34, 56, 57^. The mechanisms underlying such a bias remain an outstanding question in the recent literature on body representation ^58^. The link between body representation and motor control is a matter of great interest. From a side, the sensorimotor system might not critically depend on stored long-term memories of the body given that the brain is able to produce constant adjustments to motor output^59^. Otherwise, it has been proposed that body distortions are strictly related to motor use ^20^, as supported by studies after immobilization ^60, 61^ and in different experts, such as magicians ^62^ or football players ^63^. In patients with sensorimotor impairments after stroke, an asymmetry in the perceived arm length has been reported with the affected arm perceived as shorter as the postlesional one ^31, 64^.

This view has led to the hypothesis that in aging, alterations in mBR could be linked to diminished accuracy of the motor system ^65^. The two previous studies showing a shortage in the perceived arm length in older adults speculated about a possible role of relative underuse because of limited daily life necessities. Our data demonstrate that the motor component has an impact, and extend this view by indicating that multiple, not mutually exclusive sensorimotor and cognitive factors contribute to the distortions in arm perception during aging. We found that in older adults a higher error in the perception of the arm length is predicted by reduced hand strength, diminished proprioception during a dynamic task, and lower working memory. It is worth noting that these three explaining factors are not mutually correlated to each other (see Supplementary Materials). This result is in line with studies investigating mBR in young adults hypothesizing that mBR bias might be linked with a decline in peripheral mechanoreceptors at the levels of muscles and joints associated with alterations in proprioception ^14, 15, 66, 67^ or as happens in some clinical conditions due to decreased flow of sensorimotor information from/to the body ^23, 25, 31^. These considerations can also be applied to the context of aging considering that the process of muscle reinnervation cannot keep pace with that of denervation, leading to a loss of motor units. This appears to be related to a decline in neurotrophic factors concerned with motoneuron survival ^68^. Furthermore, the changes with age in muscle architecture (e.g., sarcopenia) are likely to impact muscle spindle function whose primary endings contribute both to the sense of limb position and movement ^69^. Accordingly, it has been suggested that older adults may have a reduced sense of movement as a result of a degraded dynamic sensitivity of spindle primary endings ^70^. These observations are consistent with the fact that in our model the arm grip strength in the arm has been identified as predictive of increased bias. Noteworthy, arm grip strength has been previously indicated as a crucial predictor for the identification of frailty in older individuals ^71^. Fried et al.^48^ operationalized a phenotype and definition of sensorimotor frailty in older adults, defined as a multifaceted clinical syndrome including several strength-related aspects (e.g. exhaustion, and weakness) and motor aspects (e.g. walking speed and low physical activities) linked with risk for falls, disability, hospitalization, and mortality^48^. Future, longitudinal studies might investigate whether the mBR bias observed here in healthy older adults (not frails, see Supplementary materials) has a predictive value in identifying frailty or other specific pattern of age-related decline in older adults.

Finally, related to the cognitive component, studies proposed that perceived distortions in mBR might be related to general bias of visual memory of the location of different landmarks ^18, 72^, or to a not efficient updating through the information coming from the body ^73, 74^. In our study, we found that only working memory (backward-span), and not short-term memory (forward span), significantly predicts the body perception bias. This suggests that a reduction in updating and manipulating online information is a more plausible factor contributing to mBR bias than a reduction in short-term maintenance of spatial information.

### Conclusions: proprioception, a key element of body perception in aging

Altogether, our data demonstrate a crucial role of proprioception in maintaining and updating an unambiguous percept of our own body during aging, both in terms of bias in the mBR of the arm and an increased tendency to experience ownership towards a virtual hand.

It is worth noting that with respect to other exteroceptive sensations such as vision and hearing that track mainly stimuli related to external objects, proprioception is strongly associated with one’s own body and its movements ^75–78^. Previous studies in pathological conditions indicated the loss of proprioception as a necessary, although not sufficient, condition to develop pathological alterations in BO. This is the case of patients showing dis-ownership towards the affected body part ^27, 28, 30, 31^, or claiming that another person’s limb belongs to them, i.e. Pathological Embodiment ^27, 28, 30, 31, 79, 80^. Moreover, proprioception provides information about position and limb movements and contributes to maintaining updated metric information about one’s own body parts, (mBR^59, 81^).

This might lay the groundwork to generate new prevention and stimulation strategies not only focused on sensorimotor functions but also on body perception in typical aging. This is particularly relevant considering that new methods for the rehabilitation of distortions in body representation have recently been proposed ^24, 26^ and tested in clinical populations^23, 25, 31, 64^, but not yet extended to support wellbeing in seniors.

## Methods

### Participants

Older adults. A group of 26 older participants were recruited in Switzerland or Italy (4 participants). Participants with cognitive impairments (i.e. equivalent scores < 2 for the Italian speakers^82, 83^; total score cutoff>26 for the French speakers^84^) assessed by the Montreal Cognitive Assessment (MOCA^84^), or severe depression (score >20) assessed by the Geriatric Depression Scale ^85^, or identified as fragile according to the criteria of Fried^48^ were excluded from the experiment. The final group consisted of 23 (8 male) with a mean age of 73.82 ± 6.36. The participants were right-handed as confirmed by the Flanders Handedness Survey^86^, had normal or corrected-to-normal vision, no psychiatric or neurological deficits, no pain or sensorimotor pathologies in the upper limbs, or fractures in the previous 12 months. Older participants were tested in two different experimental sessions to avoid fatigue during which all tasks were performed (VPD task; and all the sensory, cognitive, and motor tasks reported in Table 2; BL task; VPD task).

**Table 2.** Brief description of tasks exploited to assess cognitive, motor, and sensory functions in older adults.

| Assessed Function | Task Name | Task Description |
| --- | --- | --- |
| Sensory | Two Points Discrimination (tactile acuity)<sup>57</sup> | Participants were stimulated on their upper limbs (arm and hand) with two tactile stimuli of varying distances, and they reported whether they perceived one or two points. Adaptive methods were used for stimuli selection, and sensory thresholds were computed. |
|  | The midline judgment task (MJ; to evaluate visuo-spatial proprioception) <sup>43</sup> | Participants wearing VR must report when a visual sphere crossing their field of view on an arc centered on the participants' sternum is perceived as aligned with the midline of their body. |
|  | The proprioceptive Judgment task (PJ; to evaluate proprioceptive precision in passive condition) <sup>43</sup> | Participants must report the perceived position of their hidden arm passively moved in different positions localized on an arc centred on their sternum while relying only on proprioception. |
|  | The open Loop task (OL; to evaluate proprioceptive precision in dynamic condition). <sup>43</sup> | Participants wearing VR must reach different targets localized on an arc centered on the participants' sternum while their real hand is hidden from their view. |
| Motor | Speed of walk evaluation <sup>48</sup> | Taken from <sup>48</sup> criteria to define frailty. Participants were asked to walk twice (outward and return) at a normal pace. The fastest time was considered. |
|  | Physical Activity Rating <sup>48</sup> | Taken from <sup>48</sup> criteria to define frailty. Participants were asked to report their level of physical activity on a scale from 1 (min) to 6 (max). |
|  | Isometric arm grip strength <sup>48</sup> | Participants had to grip the JAMAR hydraulic dynamometer with maximum force while the forearm was bent at 90 degrees without support. Three measurements were taken and averaged. |
| Cognitive | Digit Span Forward (short-term memory) <sup>96</sup> | Participants were required to repeat a series of numbers in the same order as read aloud by the examiner |
|  | Digit Span Backward (working memory) <sup>96</sup> | To evaluate working memory. Participants were required to repeat the numbers in the reverse order of that presented by the examiner |
|  | Trail Making Test (TMT; visual attention and cognitive flexibility)<sup>97</sup> | The TMT part A requires the patient to connect circles in ascending numerical order. The TMT part B requires the patient to connect circles in ascending order, alternating between numbers and letters |

Young controls. For practical reasons, two different groups of 23 younger adults performed respectively the VPD or the BL task. Related to the VPD task, the power analyses in ^39^ indicates that a sample of 15 participants is appropriate to allow the statistical power necessary to see a correlation between the proprioceptive precisions measured through independent tasks and those fitted from the causal inference model (see below). Considering the BL task, the power analysis reported in previous studies a sample of 21 subjects is required to detect modifications in perceived dimensions of upper-limbs^57, 87^. To be conservative, we recruited 23 participants for both groups. All the participants were adults with no history of psychiatric, neurological, pain, or sensorimotor known deficits.

Specifically, for the VPD task, we recruited 23 healthy controls (12 male; mean age: 27.17 ± 4.27), while for the BL task, we recruited. a final group of 23 participants (8 male, mean age: 24.82 ± 4.09). All the participants of this study were tested in Switzerland or Italy according to the agreement given by the ethics committees CER-VD Canton de Vaud and CET Lombardia 1.

### Visual Proprioceptive Disparity (VPD) Task

#### Procedure

The same procedure reported in previous works ^39, 42^ was here used for the VPD task. During the VPD task participants sat in front of a table wearing an Oculus Rift S Virtual Reality system, with their arm resting on the table while their holding a motion controller (Oculus Touch) in their right hand. The real hand of the participants was hidden from their view while they observed a realistic hand in virtual reality moving coherently with the held controller. The task consists of a reaching movement from a starting position towards one of 7 possible targets (white spheres with 3 cm diameter) arranged at 0°, ±15°, ±30°, ±45° with respect to participants’ midline on an arc with radius equal to each participant maximum reaching distance.

The starting and target positions were fixed at the beginning of the experiment and calibrated before starting to be tailored to each participant, with the starting position close to the body and approximately at the midline of the participants’ body, on the table, and the reaching position corresponding to each participant’s maximum reaching distance. The position of the head was also calibrated in such a way that the task did not proceed unless the participants were able to see the targets with the head in a centered position.

In different trials, the spatial congruency between the position of the real hand of the participants, as signaled by proprioception, and the visual feedback from the virtual hand was manipulated, by introducing 9 possible disparities, i.e.: 0° (no disparity), ±13.3°, ±20.0° ±26.6° or ±40°. For positive disparities, the virtual hand is moved in a clockwise (CW) direction with respect to the real hand, while for negative directions the virtual hand is shifted in the counter-clockwise direction (CCW) with respect to the real hand. The task consisted of three experimental blocks. The first two blocks included the disparities 0°, ±13.3, ±26.6° or ±40° repeated for each target (7 disparities x 7targets). To include a measure of the subjective ratings of ownership at the end of each reaching movement the last block was shortened to limit the experiment duration. Accordingly, for each target, the last block included disparities equally spaced from 0°, ±20 or ±40° (5 disparities x 5 targets). Each block lasted approximately 7 minutes, and the total VPD lasted about 30 minutes.

At the beginning of each trial, while the real hand was in the starting position, the virtual hand was rotated by one of the possible disparity angles for 1 second, and the visual grey target appeared. After 1.5 seconds the target was turned green, and it represented the “go” signal for the participants who were instructed to start their reaching movement. Movement of the hand outside the starting position at any time before the “go” cue would automatically restart the trial. Participants were instructed to reach the target with their real (proprioceptive) hand and return to the starting position. During the last block (the third one), at the end of each trial, participants were requested to verbally report their subjective feeling of ownership for the virtual hand, evaluating their agreement with the statement “I felt as if the virtual hand was my hand” on a -1 (minimum agreement) to 10 (maximum agreement) Likert scale.

#### Bayesian Causal Inference Model

In the present work, we aimed to provide empirical evidence for the hypothesis that body ownership reshapes during aging. Previous works provided empirical evidence for the hypothesis that body ownership emerges from a Bayesian causal inference process ^39–41^. According to this framework, humans perceive their bodies through multiple senses, which often provide them with redundant information^24, 88^. The brain uses this redundancy as an advantage and merges the information from different sensory modalities into a robust unambiguous perception^89^. This mechanism of integration is defined as optimal because it produces not only lower but the lowest-variance estimate that can possibly be achieved by the Nervous System. Applied to our case of visuo-proprioceptive integration, and translated to mathematical terms, Bayesian probability theory states that the best way to interpret a stimulus S_VP_ is by performing a weighted estimation between the available visual S_V_ and proprioceptive S_P_ cues, where the weights should be proportional to the reliability of the cues^89^:

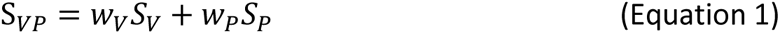

where the weights, visual weight, W_V_, and proprioceptive weight, W_P,_ sum up to unity (W_V_ + W_H_ = 1;) and should be proportional to the reliability R (i.e. the inverse of the noise of the corresponding cue 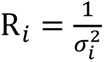) of the stimulus:

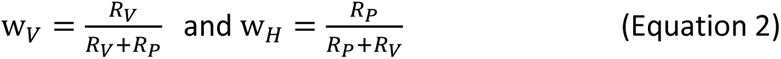

Assuming that the unimodal cues are independent, normally distributed, and originating from the same unique object of perception (i.e. forced fusion; FF), the formula that an optimal brain is expected to apply to estimate hand position is:

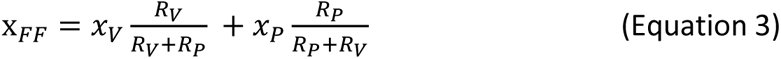

where X_FF_ is the final estimate of hand position (i.e. the one used to calibrate reaching movements in our task), and x_v_ and x_p_ are the unimodal (and noisy) position estimates. However, one of the above assumptions is non-trivial. Indeed, during everyday life situations it is not always reasonable to integrate sensory signals. For the signals to be integrated the brain has to know exactly which signals are derived from the same object or event. This is a form of correspondence problem, which has to be solved before the signals can be integrated to determine whether the object of perception is a part of one’s own body or not ^89–91^. According to Bayesian Causal Inference models, also this additional process can be described as a probabilistic event, resolved by the brain by statistically inferring the probability (P_com_) that an object is one’s own hand. Accordingly, the sensory cues involved in the process should be weighted not only according to their reliability (as in equation 3), but also considering the probability P_com_ that the final estimate corresponds to one’s own hand. Thus, the equation describing the final estimate of hand position according to the Bayesian Causal Inference (*x_BCI_*) can be refined as follows:

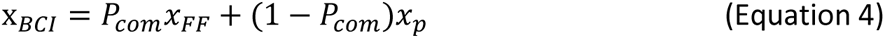

The probability P_com_ of one object (i.e.: the virtual hand) to one’s own hand varies from 1 corresponding to the certainty that the virtual hand is one’s own to 0 corresponding to the certainty that the virtual hand is not one’s own. In the model, P_com_ variations would essentially depend on the spatial disparity between its location and the proprioceptively perceived hand location:

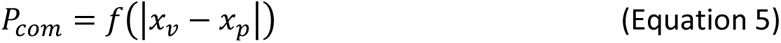

According to this function, it is clear that P_com_ depends on visuo-proprioceptive disparity (|xv-xp|). Second P_com_ is maximal at zero visuo-proprioceptive disparity and decreases as disparity increases. Third, the less vision and/or proprioception are precise, the larger the disparity will be needed by the brain to detect an incongruence. Hence, if visuo-spatial and/or proprioceptive precision is reduced, P_com_ will decrease less rapidly as disparity increases, leading to embodying the virtual hand up to larger levels of discrepancy. Fourth, the larger the prior common cause is, the larger P_com_ is at all disparities, also leading to an enhanced tendency to embody and follow the virtual hand in reaching. See Supplementary Materials for more details.

#### Bayesian Causal Inference framework transposition to Visual Proprioceptive Disparity (VPD) task

When reaching a target in the VPD task, subjects combine spatial proprioceptive information about the position of their real hand with incongruent spatial information from the virtual hand visually perceived. According to the Bayesian causal inference framework, the final reached position toward the target will be a weighted average between pure visual and pure proprioceptive guidance. The relative weight of vision and proprioception will depend on their relative precision (variabilities 𝜎_𝑉_ *and* 𝜎_𝑝_) and on the probability computed by the brain that the virtual hand is one’s own (P_comPrior_). Accordingly, if more weight is given to the visual signal (i.e. the virtual hand) than to the proprioceptive, larger reaching errors are expected in the presence of visuo-proprioceptive incongruence and/or more subjective ownership is reported towards the virtual hand (higher values of P_com_). On the opposite, pure proprioceptive guidance is considered as “zero error”, given the instructions of the task to try to reach the target with the real hand. P_com_ values can be inferred from reaching errors by computing the relative weight of vision and proprioception. P_com_ will decrease with increasing incongruence so that the relative weight attributed by subjects to the virtual hand will be high at small values of disparity and gradually decrease with incongruence. Furthermore, Equation 5 implies that if the sensory precision of vision and/or proprioception is reduced (e.g., by aging), P_com_ values will need larger disparities to decrease significantly.

To infer P_com_ σ_v_ and σ_P_ for each subject, we fitted the Bayesian causal inference model on our multisensory reaching task (VPD) using the BADS Matlab optimization tool (https://github.com/lacerbi/bads; ^92^). Following the approach used in previous studies ^39, 40^, we simulated 50000 trials for each spatial disparity. A cleaning of the data for each participant to remove systematic biases in reaching possibly due to tracking or VR calibration was done by subtracting the Observed reaching error, defined as the distance between the position of the target and the position of the real hand, the mean reaching bias at 0° spatial disparity:

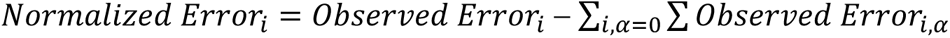

No other pre-processing step was performed on the data.

This same procedure has already been used and validated in previous studies ^39, 42^. However, we decided to further validate the model by comparing the individual parameters σ_V_ and σ_P_ extracted from the multisensory task through the Bayesian Causal Inference model with their unisensory correspondents measured through independent tasks assessing proprioception in proprioceptive static (proprioceptive judgment task; PJ) dynamic (open loop task; OL) and visual (Midline Judgment task; MJ) conditions. Previous studies have shown that the most relevant task for model validation is the Proprioceptive Judgment (PJ) which showed the best correspondence between fitted and measured parameters. However, we decided to administrate to the participants all three tasks designed to match the VPD setup, to capture the relevant unisensory components as accurately as possible in our sample. If the task relies on the proposed CI process, we expect to observe a positive correlation between the fitted parameters and at least between one of the measured unisensory parameters. Therefore, we assessed whether the parameters extracted from the unisensory tasks positively correlated with the ones extracted from the multisensory task by performing significance tests on Pearson correlation scores.

As expected, and in line with ^39^, the correlation of the σ_P_ fitted by the model with the proprioceptive precision extracted from the PJ (σ_PJ_) was significant. No correlations were found between the fitted σ_P_ and the proprioceptive precision extracted from the OL (σ_OL_) and MJ task (σ_MJ_; the detailed extraction of unisensory parameters is described below (see details on unisensory parameters extraction below section).

All the tasks have been described elsewhere ^39^. Participants were asked to sit on a chair keeping their head and trunk aligned while wearing a head-mounted display (HMD). A custom-made software programmed in Unity was used for the tasks.

The PJ task assesses the proprioceptive accuracy of the participants in static condition. During the task, the experimenter passively moved participants’ real hands to one out of 5 possible target positions selected randomly and arranged at 0°, ±10, ±20, ±40° with respect to participants’ sternum on an arc with radius equal to each participant maximum reaching distance and centred to the body midline. Participants’ real hand was hidden from their view, and at the first trial of the series, a virtual hand was displaced at +30° (right) or -30° (left). The task was a two-alternative forced-choice where the participants had to report whether the virtual hand they were observing was at the right or left of their real hidden hand. After the first trial, the position of the virtual hand was moved halving the angle and mirroring it in the opposite direction with respect to the participants’ previous answer. The sign of the initial angle was randomized trial by trial. 5 steps converging algorithm was used to determine where the participants perceived their hand to be. The final proprioceptive estimation was computed as the intermediate hand position between the last displayed position in which the algorithm converged and the next position that would have been displayed by the algorithm according to the participants’ last answer. Each target position was tested 4 times in randomized order, for a total of 140 trials. The σ_PJ_ corresponds to the variability in the perceived hand localizations along the different trials.

The OL task assesses the proprioceptive accuracy of the participants in dynamic condition (see Table 2). The procedure of this task is similar to the VPD task, that is participants were asked to do a reaching movement from a starting position to one out of five visual targets arranged at 0, ±20, ±40 with respect to participants’ sternum and administrated in random order. The targets appeared 1.5 s after the hand was in the starting position and the participants were instructed to perform the movement as soon the target turned from grey to green. Movement of the hand outside the resting position at any time during the initial resting period automatically restarts the trial. Unlike the VPD task, during the OL participants had no visual feedback (no virtual hand displayed) of their hand during the entire duration of the task. Each target position was displayed 10 times for a total of 50 trials for each participant. The σ_OL_ is computed as the variability in proprioceptive error in reaching the target.

The MJ task aims at evaluating the visuo-spatial precision of the participants. On each trial, a white sphere with a 3 cm diameter moved horizontally across the participants’ field of view at a speed of 10°/s, starting from ±45°, ±40°, ±35°, and ±30° from the body midline, on an arc centred on participants’ sternum, with a radius equal to their maximum reaching distance, as in the VPD. Participants must orally report when they felt that the visual cue was aligned with the midline of their body. The experimenter presses a button to stop the sphere, and then the visual target can be subsequently adjusted on the left or the right to match the perceived midline position. The task was repeated for 9 trials and the starting positions of the visual cue were randomized across trials. The σ_MJ_ is computed as the variability in the localization of the position in the space perceived as aligned with the body midline.

#### VPD Statistical Data Analyses

To investigate differences in the explicit ownership of the participants we fitted a linear mixed model (estimated using REML and BOBYQA optimizer in R) to predict the explicit ownership (consisting of the explicit score provided to the participants at each trial of the last block) with group and disparities (formula: ownership ∼ group * disparities). The model included ID as a random effect (formula: ∼1 | ID). The package report was used to report the models’ results according to best practice guidelines ^93^.

The model’s total explanatory power is substantial (conditional R2 = 0.69) and the part related to the fixed effects alone (marginal R2) is 0.55.

The same procedure and packages were used to investigate and report differences in the implicit ownership of the participants corresponding to the reaching error computed for each participant and each trial of the VPD task (formula: Reaching Error ∼ group * disparity). The model’s total explanatory power is substantial (conditional R2 = 0.52) and the part related to the fixed effects alone (marginal R2) is 0.38.

To demonstrate that reaching errors at the VPD can be considered a proxy of subjective ownership, we tested the hypothesis that, at the individual trial level, a higher weight attributed to the virtual hand was associated with higher subjective ownership, even at a fixed disparity. To do so, we computed the “residual” subjective ownership, by subtracting from each rating the average rating at the same disparity. Similarly, residual reaching errors were obtained by subtracting from the reaching error its average value for each disparity. To indicate a larger visual weight with positive residual drift and vice versa, residual drift values were multiplied by the sign of the spatial disparity. Zero disparity values were excluded as they yield no meaningful information in this analysis. Then, we tested whether residual error and ownership values were correlated by means of a linear mixed model.

To better understand the sensory and cognitive component underlying the ownership mechanism of the participants in the section *Factors underlying the age-related distortions in body perception: a fundamental role of proprioception* we analysed the unisensory precision extracted from the VPD (sensory: σ_v_, σ_P,_ and cognitive: P_comPrior_) and the unisensory task (σ_PJ_; see previous paragraph to have the details on the variables extractions). First, to investigate differences in the sensory precision and P_comPrior_, between groups we performed a nonpaired Wilcoxon test. Paired t-tests were performed within each group to detect differences in sensory precision between modalities. Finally, Spearman correlations tests were performed to investigate the relation between the proprioceptive precision σ_PJ_ and: i) the sensory variability extracted from the model σ_PJ_, to demonstrate that the sensory variabilities fitted from the model show correspondence with the proprioceptive variability directly observed during a proprioceptive task; ii) subjective ownership ratings, to assess the model’s validity in explaining alterations in BO through unimodal sensory deficits.

### Body Landmarks Localization Task

The body Landmarks Localization task used here has been described elsewhere ^31, 87^. To calculate the width and length of the two body parts (hand and arm), we considered the position (real and perceived) of the five landmarks (see Fig. 2A). The hand length was calculated as the mean of the distance between the tip of the annular and the external wrist and the distance between the tip of the index and the internal wrist. The arm length was obtained by calculating the mean between the other two distances, i.e. the distance between the marker on the internal wrist and the elbow and the distance between the external wrist and the elbow. The hand width was calculated as the distance between the tip of the index and annular fingers, while the arm width was obtained by calculating the distance between the internal and external parts of the wrist. Then, for each participant, we calculated an index of the bias in the perceived dimension with respect to the actual one (estimated dimension^87, 94^), as the ratio between the perceived and the real size for each body part (arm length, arm width, hand length, hand width):

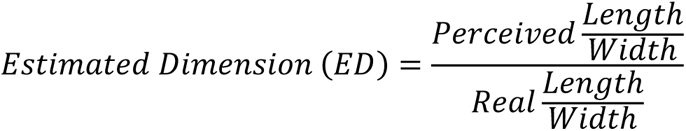

Values below 1 represent an underestimation of the perceived dimension with respect to the real one, and values above 1 indicate an overestimation. Moreover, we computed the Estimated Error index, i.e. an index representing the length/width estimation bias, independently from the underestimation or overestimation. The Estimated Error provides information related to the accuracy of the size estimation, and it is computed as follows:

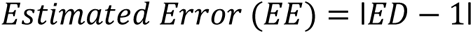

In other words, the Estimated Error is a measure of the error between the Estimated Dimension (ED) and the correct dimension (i.e. ED=1). The lowest possible value is 0 and it represents no error in the perception of the estimated length/width. On the opposite, higher values represent higher errors.

#### BL Statistical Data Analyses

To investigate bias in metric body perception we compared the performances between the young and older groups. First, we checked the normality of the distribution of the indexes using the Shapiro-Wilk test. Then we measured the distance between the Estimated Distance Index of the two groups by running an unpaired Student’s t-test for independent samples with the software R Studio (R Core Team, 2017, http://www.R-project.org/). If the distributions were not normal, we performed the analyses with the Wilcoxon signed-rank test. Bonferroni corrections were applied to correct for multiple comparisons, and it was achieved by dividing the probability value by the number of tests conducted (i.e. the number of body parts tested).

To evaluate the sensory-motor and cognitive factors underlying the bias in arm length perception observed in older adults we performed a Linear regression with the arm length Estimated Error as the dependent variable and the sensory-motor and cognitive factors associated with aging as regressors. Before running the analysis, we checked the normality distribution of the dependent variable, and we performed correlations between predictors to avoid collinearity issues. The final set of variables included in the Linear regression after checking the independence of the observations (second assumption) is reported in the results (Table 1). To find the subset of variables in the data resulting in the best performing model (i.e. model with lower prediction error) we performed a stepwise regression using stepAIC function in MASS package with R Software, the final set of variables included after the stepwise regression is reported in the Results section (Table 1).

### Sensorimotor and Cognitive Tasks’ Battery

To evaluate the sensory-motor and cognitive factors underlying the bias in arm length perception observed in older adults we asked the participants in this group to perform a battery of sensorimotor and cognitive tasks. Accordingly, we referred to a known protocol already proposed and largely used in aging to assess sensorimotor aspects declining in aging (i.e. frailty phenotype) ^48^

Further, we added a few sensory tasks to quantitively measure other potential sensory, motor, and cognitive aspects known to be associated with aging. The Tactile Estimation Task was found to be correlated with mBR ^46^. The MJ, OL, and PJ have been suggested to be relevant in body perception processing ^39^. The digit span and TMT are among the most used validated tests (not already included in the Fried protocol) performed during aging^49, 50, 95^. Table 2 summarizes all the tasks included in the battery specifying the assessed function for each task.

More details related to the PJ, MJ, and OL tasks are provided in the paragraph on *Bayesian Causal Inference framework transposition to the Visual Proprioceptive Disparity (VPD) task*.

Related to the Two Points Discrimination Tasks used to investigate the participants’ tactile acuity, we adapted the method used in ^57^. Participants were blindfolded while sitting on a chair with their right arm resting in a prone position. We measured the two-point discrimination threshold (2pdt) on the thenar eminence of the right hand, and on the center of the internal forearm along longitudinal orientations, the order of administration of the two body parts was randomized between participants. Subjects were tactilely stimulated using a caliper, and at each trial, their task was just to report whether they perceived one or two touches on the skin. The starting double posts separation was clearly above the 2pdt according to normative values ^98^, and corresponded to respectively 2 cm for the hand and 4 for the forearm. The separation was then reduced progressively by 50 % after each set of three successive correct responses. When subjects made an error, the separation was subsequently increased to the midpoint of the current (erroneous) trial and the immediately preceding (correct) trial. An increase of 150% was achieved only in the case where the starting distance (2 or 4 cm) was perceived as 1 point. This procedure was terminated after 5 inversions of the staircase. The mean between the last two distances perceived correctly was taken as an individual threshold. We then confirmed this 2pdt estimate by delivering five double posts at this separation randomly intermixed with five single posts. If subjects made less than 3 errors, the estimated threshold was accepted for experimental testing. Otherwise, the procedure was repeated.

The speed of walk evaluation, physical activity rating, and max arm grip strength evaluation were done following the Fried protocol. Related to the speed of walk evaluation, the participants were asked to walk along 4 meters distance twice. They were asked to use their habitual speed of walk. The starting and arrival points were marked on the floor and an experimenter walked behind the participant to ensure safety. The fastest walking time is considered. The Physical activity rating consists of a Likert scale from 1 (min) to 6 (max), taken from the Minnesota Leisure Time Activity questionnaire^99^. Each step of the scale suggests a level of physical activity and the participant must choose which one better describes her/his current situation. During the administration of the questionnaire, the experimenter shared some definitions by specifying that: light-intensity exercise means an activity that does not cause sweating, and that allows the person to speak; moderate intensity exercise means an activity causing sweating, and not allowing the person to speak; high-intensity exercise means maximal effort. Finally, the max strength evaluation is performed by asking the participants to sit having their right arm not supported and flexed by 90°. The subjects are asked to hold the Jamar and at the “start” signal to squeeze it as hard as they can. The procedure is repeated 3 times and the meaning of the three evaluations is considered as a measure of arm grip strength. Concerning cognitive tests, we used the Trial Making Test^100^ which is a well-known measure of cognitive flexibility^101^ in which participants were asked to connect in the correct sequence numbers (part A) and letters (part B) as fast as possible. In case of any error, the experimenter asks the subject to perform a correction, before stepping to the following sequence. The time needed to accomplish the tasks is recorded. Finally, we asked participants to perform the digit span forward and backward. The tasks traditionally are comprised in the Wechsler Adult Intelligence Scale-WAIS^102^ which is used as a general test for intelligence and has been developed to assess cognitive ability for adults. The Digit span forward requires the child to repeat numbers in the same order as read aloud by the examiner and it is considered as a good measure of simple short-term memory. The Digit Span backward represents a qualitatively different type of task. It requires the participants to repeat the numbers in the reverse order of that presented by the examiner, and it is traditionally considered as a task relying more upon working memory skills that should be considered separately from digits forward^103^. For each participant the tasks were always administered by the same experimenter and the span, i.e. the maximum number of digits that the participants were able to recall was registered.

