## Supplementary Materials for "Proprioception impacts body perception in healthy aging – insights from a Psychophysical and Computational Approach"

### Older Adults’ Sample Details

Here below we report tables detailing 1) demographic info and the scoring obtained by the participants at the tasks administered for checking the inclusion criteria, 2) the order of administration of the tasks.

Table S1. Demographic information and inclusion criteria details for the older adults’ sample.

| **ID** | **Age** | **Gender** | **Handedness** | **GDP** | **MOCA FR** | **MOCA ITA** | **Frailty** |
| --- | --- | --- | --- | --- | --- | --- | --- |
| HO_01 | 73 | M | 10 | 2 | NA | 4 | 1 |
| HO_02 | 72 | F | 10 | 11 | NA | 4 | 1 |
| HO_03 | 72 | M | 10 | 0 | 30 | NA | 0 |
| HO_04 | 81 | M | 10 | 0 | 27 | NA | 0 |
| HO_05 | 78 | F | 10 | 1 | 27 | NA | 1 |
| HO_06 | 77 | F | 10 | 4 | 27 | NA | 0 |
| HO_07 | 73 | F | 9 | 3 | 27 | NA | 1 |
| HO_08 | 76 | M | 10 | 4 | 23 | NA | 0 |
| HO_09 | 66 | F | 10 | 4 | 28 | NA | 0 |
| HO_10 | 74 | F | 10 | NA | 19 | NA | 0 |
| HO_11 | 83 | M | 10 | 6 | 27 | NA | 0 |
| HO_12 | 80 | F | 10 | 4 | 26 | NA | 2 |
| HO_13 | 65 | F | 10 | 7 | 28 | NA | 0 |
| HO_14 | 65 | F | 10 | 1 | 27 | NA | 0 |
| HO_15 | 72 | F | 10 | 6 | 28 | NA | 0 |
| HO_16 | 65 | M | 10 | 2 | 30 | NA | 0 |
| HO_17 | 66 | F | 10 | 8 | 22 | NA | 0 |
| HO_18 | 74 | F | 10 | 0 | 28 | NA | 0 |
| HO_19 | 83 | F | 10 | 2 | 29 | NA | 1 |
| HO_20 | 68 | F | 10 | 0 | 29 | NA | 1 |
| HO_21 | 75 | F | 9 | 3 | 29 | NA | 0 |
| HO_22 | 73 | F | 10 | 0 | 27 | NA | 0 |
| HO_23 | 66 | M | 10 | 9 | NA | 3 | 1 |
| HO_24 | 88 | M | 10 | 2 | NA | 4 | 1 |
| HO_25 | 74 | F | NA | 5 | NA | 4 | 2 |
| HO_26 | 75 | M | NA | 4 | NA | 4 | 1 |

The age and gender are reported for each participant.

The Handedness column reports the scoring obtained by each participant to the Flanders Handedness Survey^1^, where the participants have to rate their preference to use right or left hand in ten different situations.

The GDP column represents the scoring obtained by each participant on the Geriatric Depression Scale^2^. A score above 20 indicated severely depressed participants. None of the participants exceeded this score.

The MOCA FR, and MOCA ITA columns indicate respectively normative scoring used to assess the overall cognitive performance of the French and Italian speakers with the Montreal Cognitive Assessment test. According to the guidelines we used a total score cutoff>26 to identify cognitively healthy participants among French speakers^3,4^ and equivalent scores < 2 for the Italian speakers^5,6^. Of the twenty-six participants recruited, three were excluded because their MOCA scores did not meet the cut-off (greyed out in table S1).

Finally, the column Frialty indicates the total score of the frailty test proposed by Fried to identify frailty individuals in older adults^7^. According to the standardized ascertainment test, at least 3 of these 5 criteria must be present:

1.	Involuntary weight loss of at least 5 kg in the last 12 months or a BMI < 18.5 kg/m2

2.	Weakness, assessed by a 20% decrease in grip strength adjusted for gender and BMI

3.	Fatigue and exhaustion, self-assessed

4.	Decreased physical activity based on kilocalories expended per week (< 383 kcal for men and 270 for women)

5.	Slowness based on 4-metre walk time adjusted for gender and height

Each of these components is assessed and a maximum score of one can be assigned to participants showing difficulties in that area (i.e. they performed worst compared to the average comparison values corrected for gender and age). A sum of the scores of the 5 criteria higher than 3 indicates frailty. No participants appeared to be fragile in our sample.

Besides these tasks for checking inclusion criteria, we also add other assessments to evaluate cognitive and sensorimotor variables (namely the Digit Span Forward and Backward and the Trail Making Test for the cognitive factor, the Isometric handgrip strength for the motor factor and the Two Points Discrimination and the unisensory Proprioceptive Judgment (PJ), Open Loop (OL) and the Midline Judgment (MJ) tasks in virtual reality for the sensory factor; see Table 2 in the main text for more details), together with Body Landmark localization (BL) Task to assess the metric body perception and the Visuo-Proprioceptive Disparity Task (VPD) to assess the ownership experience. All the tasks were administered in randomized order usually within two (max 3) sessions, according to participants availability and fatiguability. The sessions must be separated at least 30 minutes one from the other. The Body Landmark Localization Task (BL) was performed before the VPD, to avoid the movement and observation of the arm for the entire duration of the VPD affecting the BL results.

Concerning the VPD, in line with the previous studies ^8,9^, the unisensory tasks: the PJ, the Open Loop task OL, and MJ, were always administered before the VPD. The cognitive and sensorimotor tasks were administered in randomized order.

### Independence of regression model variables

The first assumption to perform linear regression is that the observations are independent. Accordingly, we performed Spearman correlation tests to check that the predictive variables that were assessed to predict the bias (see Table 1 in the main text) were not significant correlated one with the others.

We found a significant correlation between visual attention assessed measuring the time needed to perform the Trail Making Task (TMT) A and both the speed of walk [r(21) = 0.45, p =.03] and the proprioceptive variability in dynamic condition [r(21) = 0.44, p =.03], while these two last variables did not correlate one with the other. The tactile acuity on the hand (2PD) was significantly correlated with the tactile acuity on the arm [r(21) = 0.42, p =.04] and the proprioceptive variability in static condition assessed with the PJ task [r(21) = -0.46, p =.03], while these two latter variables were not correlating each other. Finally, the visuo-spatial variability assessed with the midline judgment task (σ_MJ_) was correlated with the proprioceptive passive variability (σ_PJ_) [r(21) = 0.47, p =.03]. All the parameters that did not correlate one with the other were kept in the analyses.

Accordingly, the time to perform the TMT-A, the tactile acuity on the hand and the visuo-spatial proprioceptive variabilities (σ_MJ_) were removed to ensure no dependency between variables included in the regression model by removing the lowest possible number of variables.

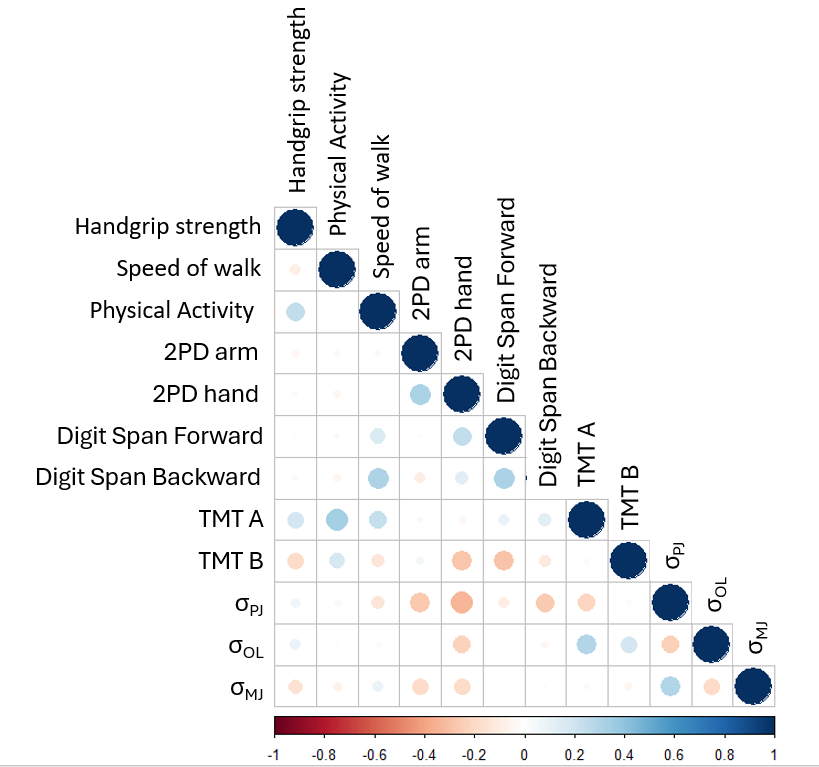

Figure S2. Correlation Matrix between regression model predictive variables. Each row represents the correlation between each variable and the others. The colors represent positive (in blue) or negative direction of correlation, the color intensity represents the strength of correlation.

### Correlation between proprioceptive and visual variability extracted from the model and those obtained from unisensory tasks

In order to validate whether the unisensory variability extracted from the model gave a good measure of the visual and proprioceptive abilities of the participants (and according to previous studies exploiting the same approach ^8^), we decided to introduce dedicated tasks, namely the Proprioceptive Judgment (PJ), the Open Loop (OL) and the Midline Judgment (MJ) (see description of the task in the Method section of the main text). The selection procedure of the type of tasks to include is detailed in ^8^, and several options were considered. Overall, the PJ was selected as a measure to assess proprioception under static conditions. Given that the model from the multisensory task incorporates motor components that might be not captured by a purely proprioceptive task, in addition to the PJ, an open-loop reaching task (OL) was used to isolate the combination of motor and proprioceptive components which can determine the end-point precision in a reaching task. Concerning visual precision, the visual midline judgement (MJ) task was used as the closest approximation to isolate the contribution of visual information to position estimates in our task. These tasks allowed us to measure proprioception and vision independently from the main multisensory tasks. Accordingly, to validate the parameters extracted from the model we assessed the correlations of the variabilities extracted from the model (visual: σ_V_, proprioceptive: σ_P_) with the corresponding variabilities extracted from unisensory tasks (σ_PJ_, σ_OL_, σ_MJ_). As reported in the main text we found a significant correlation between the observed σ_PJ_ and extracted σ_P_ variabilities [r(44) = 0.554, p < .001]. This result is in line with previous findings^8,9^ and suggests that the prediction of the model well describe the observed ability of the patient when considering a static proprioceptive task. In contrast, as visible in figure S2 neither the measured visual (σ_MJ_) [r(44) = 0.24, p =.21] nor the σ_OL_ [r(44) = 0.18, p = .23] precisions correlated with the corresponding unisensory variabilities extracted from the model (respectively σ_V_, σ_P_ ) suggesting that in this study these two abilities do not directly represents the sensory abilities underlying the multisensory reaching VPD. Furthermore, we found no significant differences in the σ_MJ_ [Wilcoxon Signed-Ranks Test: z = -0.610, p=.542] and σ_OL_ [z = -0.741, p=.459] of the two groups of young and older adults.

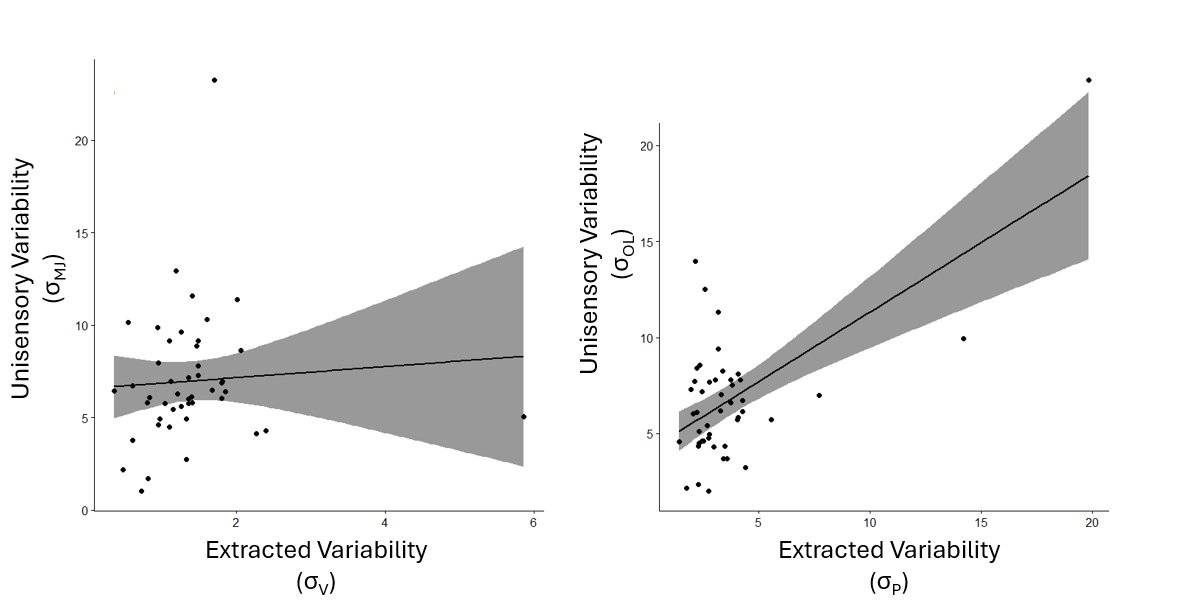

Figure S1 Relation between extracted and observed visual (left panel) and proprioceptive (right panel) variabilities.

### Disownership of the real limb in elderly

In a previous work on stroke patients, we found that the increased tendency to embody an incongruent virtual hand of the patients at the VPD task was related to the disembodiment of patients’ real hand ^9^. In other words, we found that patients not only reported higher implicit and explicit ownership towards the virtual hand when there was a disparity with their real hand, but also, they reported less ownership towards the virtual hand when it was actually congruent with the real hand, i.e. at zero disparity. To investigate whether this was the case also for the older adults in this study we assessed if the proprioceptive variability, that we demonstrated being at the basis of tendency to embody the incongruent virtual hand, is also explaining a disownership at zero disparity. To do that, we performed Spearman correlation between the measured proprioceptive variability assessed with the PJ task and both the Reaching Error and the Explicit Ownership Ratings measured at the VPD task in condition of 0 disparity. We found no significant correlation either between the σ_PJ_ and the Explicit Ownership Ratings [r(44) = 0.10, p =.51] nor the Reaching Error [r(44) = -0.02, p = .89].

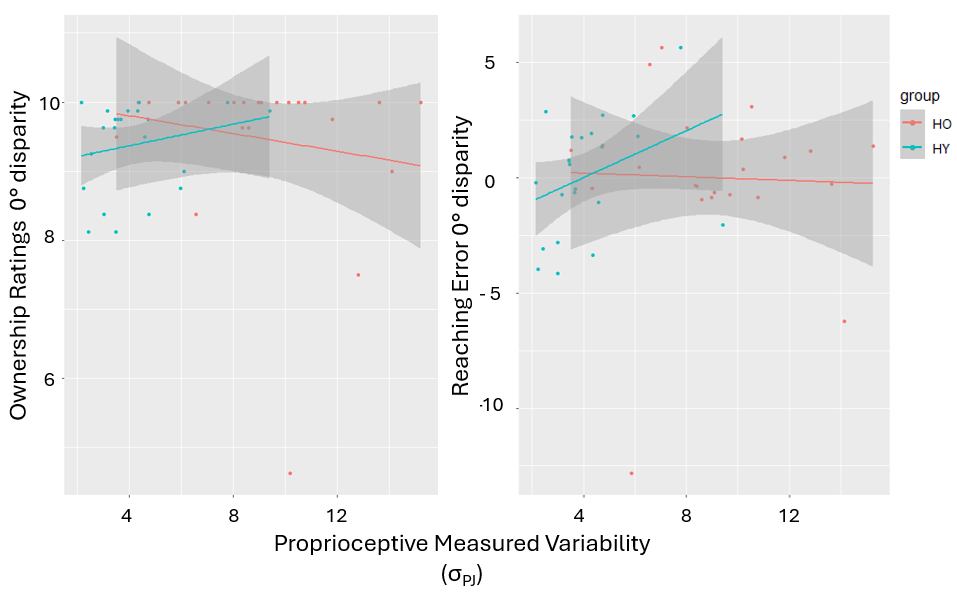

Figure S4. Relation between Proprioceptive Variability (σ_PJ_) assessed through the Proprioceptive Judgment task and the Explicit Ownership Ratings (left panel) and Implicit Reaching Error (right panel) reported at 0° disparity condition (i.e. when the virtual and real hand where in congruent positions).

We found no evidence of disownership of the real limb in the older participants, even when making inter-group comparisons, neither by comparing the explicit ownership judgements (Wilcoxon signed-rank: z -2.47, p=.013), nor by considering the reaching error (Wilcoxon signed-rank: z = -0.13, p=.896).
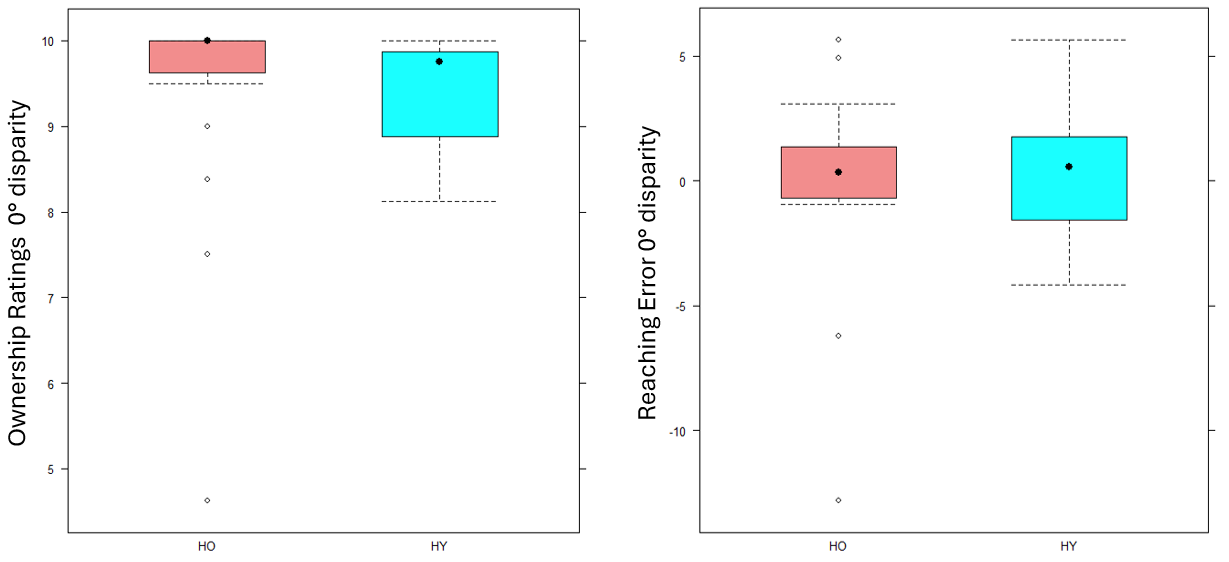

Figure S5. Comparison between the Explicit Ownership Ratings (left panel) and Implicit Reaching Error (right panel) of the older (in magenta) vs the younger (in blue) adults that have been observed at 0° disparity condition (i.e. when the virtual and real hand where in congruent positions).

### Bayesian Causal Inference Framework

Following the approach used in previous studies ^8,9^, we modeled the emergence of body ownership from visuo-proprioceptive integration in a Bayesian Causal Inference (Bayesian CI) framework, that we applied to our VPD reaching task. To obtain the equations of the Bayesian CI model, we start from the sensory component, by modeling the sensory stimuli underlying those congruencies, i.e. the joint probability distribution of physical stimuli and their associated neural representation. The relevant physical variables considered in the VPD task, consist of the true positions of the real s_p_ and of the virtual s_v_ hand (expressed in degrees from the shoulder). At each trial, the virtual hand seen by the participant might be considered as “being” or “not being” the participant’s hand. In Bayesian framework it means that s_v_ and s_p_ may have one same physical cause (C=1) or two different (C=2). Where C is drawn from a Bernoulli distribution with probability P_π_:

$$P\left( C=1 \right)=P_{\pi} \left( 1 \right)$$

If C=1, then s_v_=s_p_=s meaning that the visual and proprioceptive position of the hand is the same, and is drawn from a uniform distribution on the -90/90 degrees range, approximating the set of reachable angles. If C=2, s_v_, s_p_ are drawn independently from the same uniform distributions.

In order to simulate variability in sensory inputs, Zero mean Gaussian noise was added to the sensory inputs true positions, to model the neural noise in sensory inputs encoding. That is, the internal representations of s_v_ and s_p_ become respectively x_v_ = s_v_ + N(0, σ_v_), x_p_ = s_p_ + N(0, σ_p_), and σ_v_ and σ_p_ therefore represent the visual and proprioceptive information uncertainty about hand position.

Given this information, the Bayes theorem allows to compute the posterior distributions for the positions of the stimuli and the number of underlying causes that an ideal observer would compute. So, starting from the number of causes of the observed stimuli, we have:

$$P_{\mathrm{com}}=P\left( C=1\vee x_{p},x_{v} \right)=\frac{P\left( x_{p},x_{v}\vee C=1 \right)P\left( C=1 \right)}{P\left( x_{p},x_{v}\vee C=1 \right)P\left( C=1 \right)+P\left( x_{p},x_{v}\vee C=2 \right)P\left( C=2 \right)} \left( 2 \right)$$

In our experimental setup, P_com_ translates to the probability that the virtual hand is one’s own, where higher probabilities correspond to higher levels of subjective ownership. Accordingly, assuming that the region outside the -90 to 90 degrees range contributes negligibly to the integral, the likelihood functions defined by our generative model are:

$$P\left( x_{p},x_{v}\vee C=1 \right)=\frac{1}{\alpha}\exp\left[ -\frac{1}{2}\frac{\left( x_{v}-x_{p} \right)^{2}}{\sigma_{p}^{2}+\sigma_{v}^{2}} \right]≝\frac{1}{\alpha}e^{\frac{-\delta_{s}^{2}}{2\sigma_{s}^{2}}} \left( 3 \right)$$

$$\alpha=\iint_{-90}^{90} P\left( x_{p},x_{v}\vee C=1 \right)\approx180\sqrt{{2\pi(\sigma}_{p}^{2}+\sigma_{v}^{2})}\left( 4 \right)$$

Where α is the normalization constant, δ_s_ denotes the spatial visuo-proprioceptive disparity, and σ^2^_s_ is a short form for the sum of the uncertainties on visual and proprioceptive positions. When there are two separate causes, we simply have:

$$P\left( x_{p},x_{v}\vee C=2 \right)=\frac{1}{{180}^{2}}\left( 5 \right)$$

Therefore, the probability of common cause is given by:

$$P_{\mathrm{com}}=P\left( C=1\vee x_{p},x_{v} \right)=\frac{P_{\pi}\frac{1}{\alpha}e^{\frac{-\delta_{s}^{2}}{2\sigma_{s}^{2}}}}{P_{\pi}\frac{1}{\alpha}e^{\frac{-\delta_{s}^{2}}{2\sigma_{s}^{2}}}+(1-P_{\pi})\frac{1}{{180}^{2}}} (6)$$

The final estimate of hand position Ŝ_p_, is obtained using the same inference process. Indeed, if P_com_ = 0, visual information should be ignored, and Ŝ_p_ should reduce to the pure proprioceptive estimate. If P_com_ = 1, visual and proprioceptive information should be integrated, and Ŝ_p_ should correspond to the “classical” forced fusion estimate ^10^ in which inputs are weighted by their inverse precisions. In all the other general cases of “intermediate” P_com_, Ŝ_p_ should then be the average of the proprioceptive and forced fusion estimates, weighted by P(C=0) and P(C=1) respectively.

$$Ŝ_{p}=P\left( C=1\vee x_{p},x_{v} \right)\frac{\sigma_{p}^{2}x_{v}+\sigma_{v}^{2}x_{p}}{\sigma_{p}^{2}{+\sigma}_{v}^{2}}+P\left( C=2\vee x_{p},x_{v} \right)x_{p} \left( 7 \right)$$

In our framework, Ŝ_p_ values should be used when computing motor commands to reach a target in the presence of visuo-proprioceptive disparity, so the reaching error will be inversely proportional to the difference between the true hand position, S_p_, and its estimate Ŝ_p_. Then, equation (7) can be used to formulate behavioural predictions given arbitrary true positions of the stimuli (s_v_, s_p_), and individual values of sensory precisions and common cause prior (σ_v,_ σ_p,_ P_π_). To do so, noisy neural representations x_v_ and x_p_ are generated based on the selected values of s_v_ and s_p_, and inserted in equation (7) to compute Ŝ_p_. This process corresponds to simulating one behavioural trial.

9.	Mastria, G. *et al.* Body ownership alterations in stroke emerge from reduced proprioceptive precision and damage to the fronto-parietal network. *Rev.*

10.	Ernst, M. O. & Banks, M. S. Humans integrate visual and haptic information in a statistically optimal fashion. *Nature* **415**, 429–433 (2002).
